# Environmental life cycle assessment of recombinant growth factor production for cultivated meat applications

**DOI:** 10.1101/2023.06.01.543245

**Authors:** Kirsten Trinidad, Reina Ashizawa, Amin Nikkhah, Cameron Semper, Christian Casolaro, David L. Kaplan, Alexei Savchenko, Nicole Tichenor Blackstone

## Abstract

Growth factors are critical components of current serum-supplemented and serum-free media formulations for cultivated meat production. However, growth factors have been excluded, estimated using proxies, or modeled using proprietary data in existing environmental assessments of cultivated meat products. Cell culture media has been identified as a hotspot in such studies, therefore it is important to accurately quantify the environmental impacts of growth factor supplementation. To address this gap, this study applied life cycle assessment (LCA) methodology to comparatively assess the environmental impacts of recombinant growth factor production for cultivated meat applications. Life cycle inventories were developed for four recombinant growth factors (IGF-1, FGF, TGF-ß, and PDGF) produced using a novel bench-scale process. The functional unit of the product output was selected as 1 mg of produced growth factor. The results indicate that recombinant growth factors can have significant environmental impacts within cultivated meat systems, despite being used in very small quantities. For example, the global warming potential of production of 1 mg of IGF-1, FGF, TGF-ß, and PDGF was estimated to be 0.1, 0.04, 0.2 and 0.2 kg CO_2_ eq, respectively. Future research should explore the sustainability of producing these growth factors at scale to meet the needs of the expanding cultivated meat industry or identifying alternatives to these growth factors that have a lower impact on the environment.

**Nomenclature:** 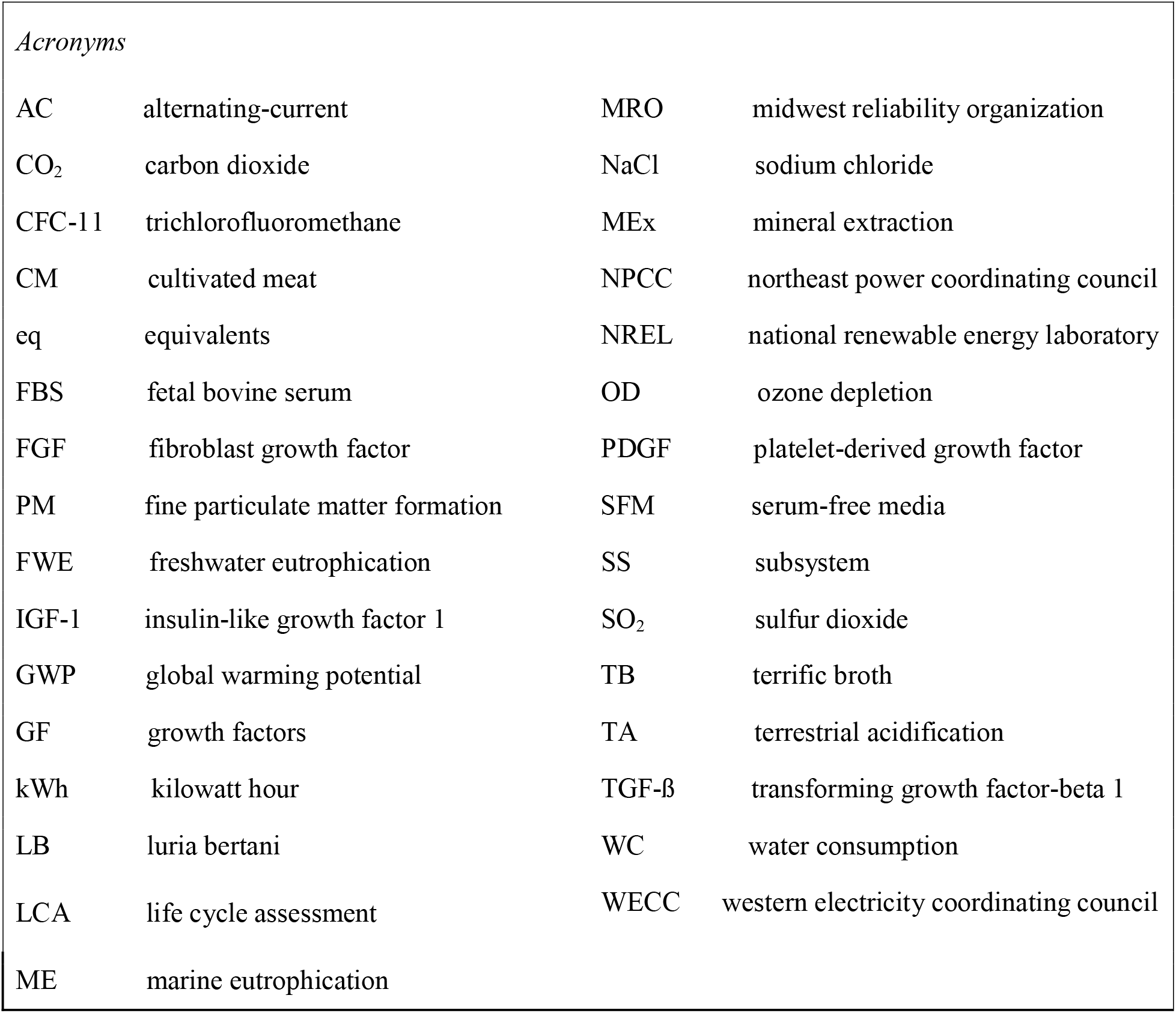

## 1. Introduction

The significant climate change, animal welfare, and human health consequences attributed to industrialized livestock production point to the need for transformative changes to the current food system (Godfray et al., 2018; Stephens et al., 2018). Cultivated meat (CM) has gained recognition as an alternative method of meat production with the potential to ameliorate these concerns (Datar et al., 2009; Mattick et al., 2015; Sinke and Odegard, 2021). CM is a subset of cellular agriculture and is produced through the *in vitro* culture of animal cells in nutrient-rich media; also termed tissue engineering.

To ensure the commercial viability of CM, cell culture media (hereafter, media) needs to be both environmentally sustainable and free of animal products (Specht, 2020). A key ingredient of conventionally formulated media is serum, which consists of an amalgam of nutrients, proteins, and growth factors (GFs) (Verma et al., 2020). Serum is most commonly sourced from the blood of bovine fetuses and processed to develop fetal bovine serum (FBS). Due to the animal-based origin and ethical concerns regarding FBS, achieving effective serum-free media (SFM) formulations has been a critical point of interest in the development of CM technology. In these formulations, recombinant GFs such as insulin-like growth factor 1 (IGF-1) have been used as protein supplements to supplant serum and promote cell proliferation and development activities (Lee et al., 2022). These supplements are a critical component to SFM as they mimic the GF composition of traditional serum-supplemented media (Venkatesan et al., 2022).

Despite their importance, published information regarding the environmental impact of GF production is either proprietary or based on pharmaceutical-grade recombinant protein production proxies. In the peer-reviewed LCA literature on CM (Tuomisto & Teixeira de Mattos, 2011; Tuomisto et al., 2014; Mattick et al., 2015; Smetana et al., 2015), GF production was left outside of the system boundary. To our knowledge, the GF production process has only been included within the system bounds of a non-peer-reviewed report released by CE Delft and a recently published CM bioprocess LCA (Sinke and Odegard, 2021; Tuomisto et al., 2022). In both papers, however, GF production was approximated using amino acid inventory proxies. While precision fermentation processes follow similar production processes, different products necessitate different parameters (i.e., production time, feedstock conversion efficacy, downstream processes). Although the CE Delft report uses primary water and electricity data from recombinant protein fermentation processes, the data for this study was retrieved from private industry — inventory and parameter data were not made publicly available. In spite of proxy usage, the CE Delft LCA reports that recombinant proteins, despite their low concentrations (<1 g/L), have high significant contributions to the overall environmental impact of media. In their review of CM LCAs, Rodriguez Escobar and colleagues (2021) identified the omission of GFs as a major limitation, highlighting that minor changes in GF usage could have considerable influence on the environmental impacts of the CM system as a whole.

Few other peer-reviewed recombinant protein LCAs have been published to date, however varying data sources, production systems, and their anticipatory nature result in highly variable results (Feijoo et al., 2017; Järviö et al., 2021). Feijoo et al. (2017) report a climate change impact of 16,755 kg CO_2_ eq per kg of recombinant ß-galactosidase protein, while an LCA performed on recombinant ovalbumin found an associated climate change impact of 9.6 kg CO_2_ eq per kg of the protein (Järviö et al., 2021). Differences in the molecular weight of the products and host organisms used are possible causes of the discrepancies in processes. Environmental impacts may also vary due to differences in processing requirements dependent on the product grade. Stricter purity requirements exist for pharmaceutical products, often requiring more intensive downstream processing steps and therefore increasing water, energy, and material usage (Puetz & Wurm, 2021). These findings indicate that, rather than relying on proxy data, modeling of product-specific recombinant protein production processes is necessary to accurately represent such a hotspot in the CM product system. To address this gap, this study uses life cycle assessment to develop inventory data and assess the environmental performance of recombinant GF production used for CM.

## 2. Materials and Methods

Life cycle assessment (LCA) is a widely used, standardized method to estimate the potential environmental impacts of products or services along their life cycles. There are four phases of LCA according to the International Standards Organization (ISO) LCA standards: goal and scope definition, inventory analysis, impact assessment, and interpretation (ISO, 2006a; ISO 2006b).

### 2.1 Goal and scope definition

The goal of this study was to construct bench-scale life cycle inventories (LCI) of CM-specific GFs using primary data and to benchmark the life cycle environmental impacts of the recombinant GF production process for CM. Commonly used in cell culture media to support muscle cell proliferation, insulin-like growth factor (IGF-1) was chosen as the focus for this work. Essential 8 stem-cell medium, a widely used and commercially available xeno-free medium used within serum-free formulations for CM-relevant cells, includes insulin at the highest concentration (Humbird, 2021; Specht, 2020; Stout, 2022). Three other GF families, basic fibroblast growth factor (FGF), transforming growth factor beta 1 (TGF-ß), and platelet-derived growth factor (PDGF), were chosen as comparators to model based on their use as Essential 8 medium components, as well as availability of primary data. Process and input data for bench-scale GF production provided by a collaborating laboratory at the University of Calgary were used to construct GF LCIs (Venkatesan et al., 2022).

This study aims to inform researchers on the potential impacts of recombinant GFs as they are an essential component of media for CM production. Further, this study serves to establish the first source of publicly-available process data for recombinant GF production. By making the inventories developed in this study available, researchers in the field will be provided with high-quality data on this critical component of the cell culture process to be used in future LCA work.

The functional unit (FU) of this study to which all input and output values were normalized was 1 mg GF, consistent with experimentally-validated lab-scale production outputs and CM media requirements for CM research and development or pilot-scale operations (Venkatesan et al., 2022; Sinke et al., 2023). The system boundary includes processes starting with the proliferation of IGF-1-producing *Escherichia coli* (*E. coli*) cells and ending after the purification of the desired GF product (Figure 1). The strain engineering and transformation processes occurring before cell proliferation, which are part of research and development, were excluded to ensure consistency with the literature (Feijoo et al., 2017; JärviöL et al., 2021; Gusarov et al., 2014; Kim et al., 2009). According to experiments conducted by the collaborating laboratory, the GFs were biologically active and soluble following dialysis and purification, requiring no additional post-translational modifications due to the use of optimal E. coli strains and fusion partners (Venkatesan et al., 2022).

**Figure 1.**
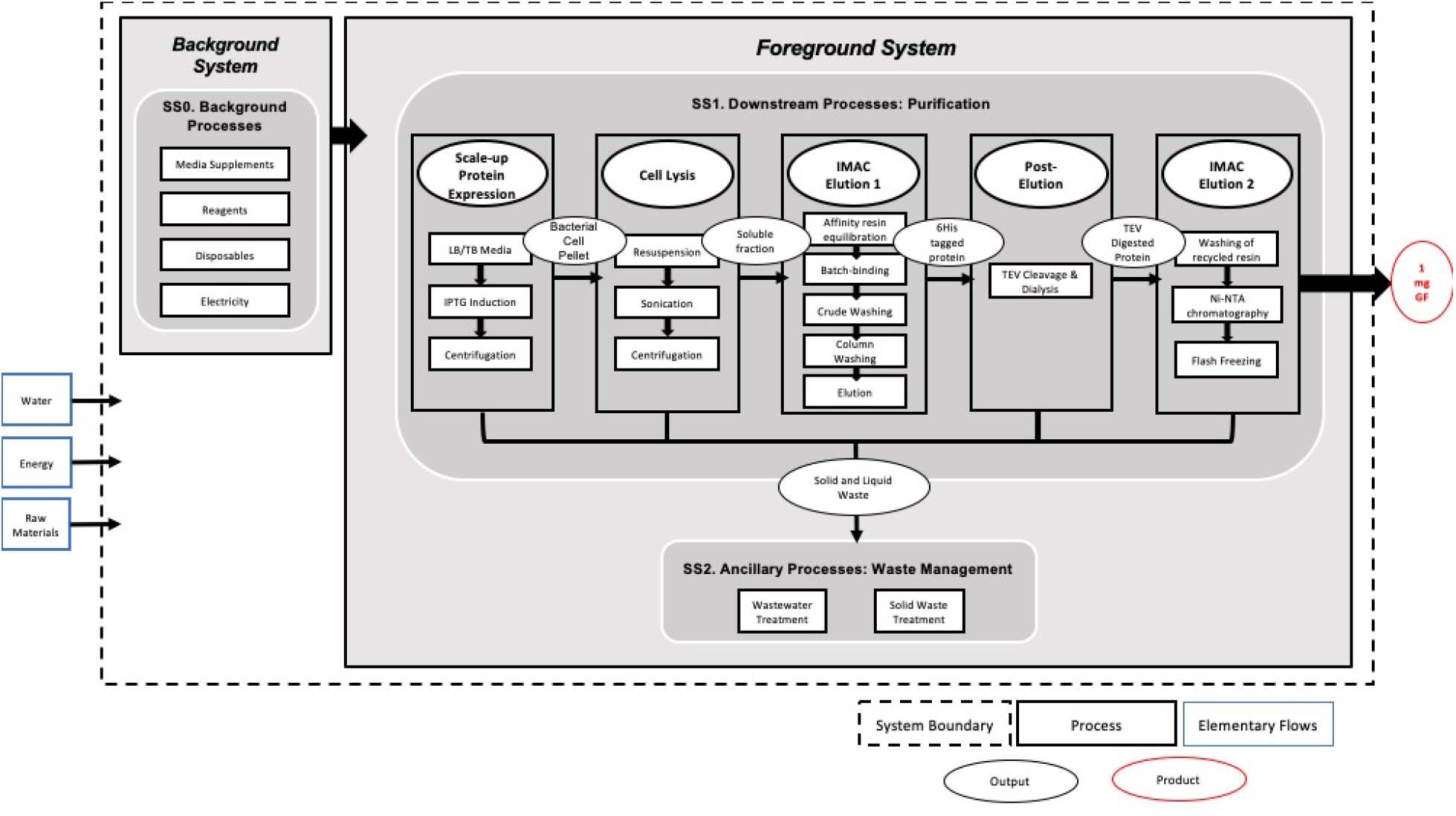
Product system diagram for bench-scale production of 1 mg IGF-1. Inputs and outputs relevant to the focus of this study are included within the system boundary, indicated by the dotted border, and normalized according to the system’s functional unit, indicated in red. The main processes comprising recombinant growth factor production are found in subsystem 1 (SS1) of the foreground system, with management processes for SS1 waste comprising subsystem 2 (SS2). Process development for SS1 inputs is captured by the background system. Resources extracted from the environment that compose the system’s inputs are represented by elementary flows in blue.

Analysis of the GF product systems was carried out using openLCA v.1.11.0 software and the ecoinvent 3.5 database, Impact assessment was conducted using the ReCiPe Midpoint 2016 (H) method and Cumulative Energy Demand (CED). The following impact categories within ReCiPe were of focus due to their importance in food systems: global warming potential (GWP) (kg CO_2_ eq), water consumption (WC) (m^3^), freshwater and marine eutrophication potential (FWE, ME) (kg P eq & kg N eq, respectively), stratospheric ozone depletion (SOD) (kg CFC11 eq), terrestrial acidification (TA) (kg SO_2_ eq), and fine particulate matter formation (PM) (kg PM2.5 eq). These impact categories address emissions and resource use along the life cycle of the product and therefore have strong relation to flows going into the environment (Huijbregts et al., 2016). Because these GFs are intended for use in food manufacturing and involve industrial production processes, categories associated with electricity and water usage were included in this study. The inventory of emissions to air, soil, and water in the production of recombinant GFs is provided in Supplementary Material 1. Coproducts are not produced by foreground processes, therefore allocation procedures were not needed. Allocation in background data (ecoinvent version 3.5) was handled according to the cut-off system model, which assigns burdens according to economic value. Cut-off criteria were used to exclude flows that had neither exact matches in the ecoinvent database nor accessible data to model as a background process. Further information can be found in Supplementary Material 3.

### 2.2 Bench-scale process descriptions

Bench-scale CM GF production includes upstream processes involving the culture and induction of transformed host microorganisms and downstream processes involving the multi-step purification of the GF proteins. Figure 1 provides an overview of the product system and system boundaries considered for bench-scale GF production, separated in terms of background and foreground systems. The background system includes subsystem 0 (SS0) and models inputs necessary for the foreground system using inventory data from the literature, as they represent novel processes not readily available in ecoinvent. The foreground system is separated into two subsystems: subsystem 1 (SS1) comprises the proliferation of *E. coli* for recombinant protein production, cell lysis, and purification, while subsystem 2 (SS2) comprises the ancillary waste management processes, including disposable and wastewater treatment. Table 1 provides a summary of input data regarding reagents, disposables, water, and energy use in the foreground system for IGF-1. Summaries of input data for FGF, TGF-ß, and PDGF-BB, as well as the input data for each phase of the GF production process are provided in Supplementary Material 1.

**Table 1.** Inventory summary of bench-scale IGF-1 product system (FU=1 mg IGF-1)

| Inputs for 1 mg of IGF-1 |  |
| --- | --- |
| *Reagents* |  |
| Tryptone | 1.1 g |
| Yeast extract | 0.56 g |
| HEPES | 0.85 g |
| Imidazole | 0.047 g |
| Sodium chloride | 2.4 g |
| Glycerol | 4.7 g |
| EDTA | 0.00039 g |
| Liquid nitrogen | 1.8 mL |
| Ethanol (95%) | 0.053 mL |
| Bleach (5%) | 1.1 mL |
| Resin | 0.059 g |
| Pierce Coomassie Plus Reagent | 0.22 mL |
| Enzymes | 0.00010 g |
| Viscose fiber | 0.033 g |
| *Laboratory Disposables* |  |
| Polyethylene | 0.87 g |
| Polypropylene | 8.4 g |
| *Water* |  |
| Purified water | 0.11 L |
| Tap water | 0.39 L |
| *Energy* |  |
| Incubation | 0.034 kWh |
| Mixing | 0.037 kWh |
| Lysing | 0.0092 kWh |
| Centrifugation | 0.013 kWh |
| Refrigeration/freezing | 0.00017 kWh |
| *Waste Treatment* |  |
| Wastewater | -0.45 L |
| Waste plastic | -9.2 g |

Recombinant GF production is modeled as aerobic fermentation of *E. coli* to reflect bench-scale GF production methods developed by the collaborating laboratory (Venkatesan et al., 2022). Transformed *E. coli* expressing a N-terminal polyhistidine (6-His) tag on the GFs were cultured overnight in Luria Bertani (LB) and terrific broth (TB) in shaking incubators at 37°C. LB/TB broth is commonly used in microbial cultures as they are both nutritionally rich media, containing yeast extract at 5 g/L and tryptone and sodium chloride both at 10 g/L. To initiate expression of the GF-producing gene within the *E. coli*, 0.6 mM of Isopropyl-β-D-Thiogalactopyranoside (IPTG) was added. The culture was then incubated overnight and centrifuged at 7,000g. Following induction and incubation, the GF-containing cells were resuspended in a binding buffer solution (pH 7.5) consisting of 100mM 4-(2-hydroxyethyl)-1- piperazineethanesulfonic acid (HEPES), 500 mM NaCl, 5mM imidazole, and 5% glycerol (v/v). Since *E. coli* cells produce the GF protein intracellularly, the cells were lysed using a sonicator and centrifuged at 20,000g to extract the GF and remove cell debris.

To purify the GF, several separation and purification steps were necessary. The first of these steps was immobilized metal affinity chromatography (IMAC), which utilized a nickel-nitrilotriacetic acid (Ni-NTA) resin matrix. The affinity column was first equilibrated with a binding buffer containing 100 mM HEPES, 500 mM NaCl, 5 mM imidazole, and 5% glycerol (v/v). When passed through the column, the GF bound to the matrix through its 6-His tag.. After binding, the column was washed using a buffer (pH 7.5) consisting of 100 mM HEPES, 500 mM NaCl, 30 mM imidazole, and 5% glycerol (v/v). The immobilized IGF-1 was finally eluted with a buffer of 100 mM HEPES, 500 mM NaCl, 250 mM imidazole, and 5% glycerol (v/v). Pierce Coomassie Plus Reagent was used at this step to track the protein during elution by mixing 200 µL of reagent with 10 µL of eluate. EDTA was added to a final concentration of 0.5 mM after elution.

The eluted product was filtered, and the 6-His tag was cleaved with 900 μg TEV protease in100mM HEPES, 500mM NaCl, and 5% glycerol (v/v), and dialyzed using PBS containing no imidazole (pH 7.4). A second affinity chromatography purification process was performed to obtain a pure GF product that was subsequently flash-frozen for storage.

### 2.3 Inventory analysis

The list of processes used for GF production LCI development can be found in Supplementary Material 2. Inputs for amino acid production background processes were modeled using data from Mattick et al. (2015) and Marinussen and Kool (2010). Additional information on key ingredients, such as induction media and equipment specifications used for energy requirements were collected from respective equipment vendors. Proxies were used when exact- or near-matches were unavailable in the ecoinvent database. Cut-off criteria were employed to exclude three reagents used in the original procedure (IPTG, benzamidine hydrochloride, and PMSF) from the modeled product system. Reagents were excluded if they lacked representative providers in available LCI databases and if they contributed less than 1% of the corresponding unit process’ total input mass. The list of proxies and excluded reagents can be found in Supplementary Material 3 alongside assumptions made for data collection and analysis.

Electricity use was estimated based on reported duration and wattage of equipment used within the laboratory processes. Because the West Coast of the United States is a hub for CM production, the Western Electricity Coordinating Council (WECC) grid mix in ecoinvent was used as a baseline, while other grids were explored to analyze sensitivity of the results (see Section 3.8). Additional information on laboratory equipment can be found in Supplementary Material 1. Capital goods were excluded from this study to be consistent with the existing literature (Mattick et al., 2015).

### 2.4. Sensitivity analyses and scenario modeling

GF production was modeled based on existing literature and primary bench-scale data. Data uncertainties in the product system necessitated sensitivity analysis, which was completed using openLCA v.1.11.0 software. Uniform distribution within a +/– 10% margin of the most relevant inputs identified within the model and in existing LCA literature on recombinant protein production were added to the product systems as parameters to be used for sensitivity analysis. Four inputs were chosen for sensitivity analysis: electricity, glycerol, disposables, and water.

Following sensitivity analysis, three scenarios were developed to explore the impact of alternative electricity grids and process designs. Two electricity scenarios were developed to compare the current WECC electricity grid mix with two alternative grid mixes from regions in the US available in ecoinvent— Midwest Reliability Organization (MRO) and Northeast Power Coordinating Council (NPCC). One scenario was developed to model an optimized version of the process, which would eliminate the optional TEV cleavage and second chromatography steps. The fusion protein acquired from this process was functionally comparable to the cleaved protein acquired after post-elution in cell growth studies.

## 3. Results and Discussion

### 3.1 Impact of growth factor production

Figure 2 shows the environmental contributions of the four GF production systems: IGF- 1, FGF, TGF-ß, and PDGF. The four GFs were produced through the same streamlined bench-scale process, with differences in impacts explained by differing product yields. The production process of FGF yields the greatest amount of purified protein, resulting in the most minimal environmental impacts, while lower yields from the PDGF production process results in higher impacts across all categories. As IGF-1 is the most commonly used GF used in media for CM applications (Humbird, 2020), the GWP and water consumption impacts specific to this GF were analyzed further in terms of sub-processes and input types.

**Figure 2.**
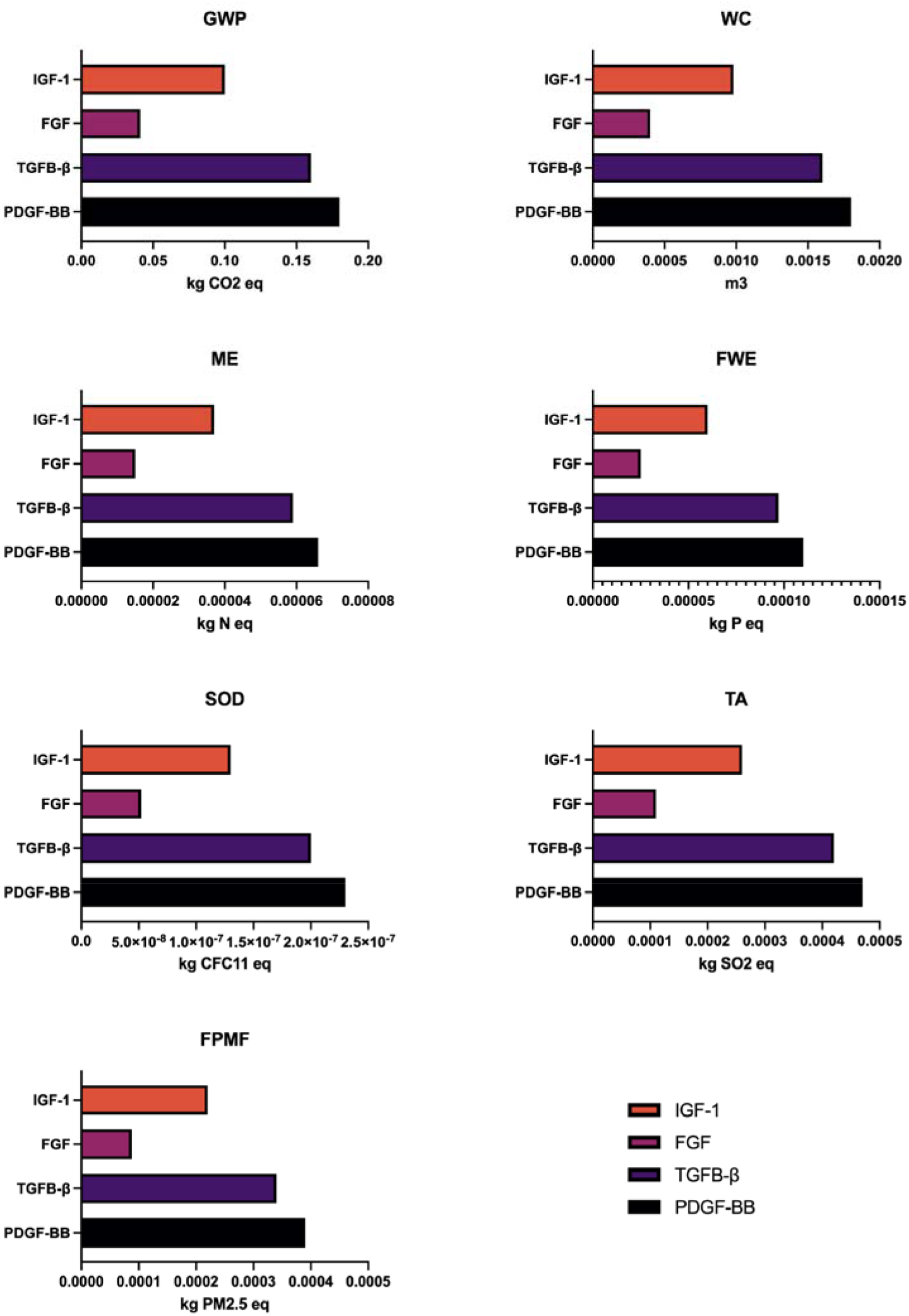
Environmental impacts of GF production per mg. Results per mg of IGF-1, FGF, TGF-ß, and PDGF products assuming identical processes with the following respective yields: 9mg, 22mg, 6mg, 5mg. Categories include: GWP– Global Warming Potential (kg CO_2_ eq), WC– Water Consumption (m^3^), ME– Marine Eutrophication (kg N eq), FWE– Freshwater Ecotoxicity (kg P eq), SOD– Stratospheric Ozone Depletion (kg CFC11 eq), TA– Terrestrial Acidification (kg SO2 eq), FPMF– Fine Particulate Matter Formation (kg PM2.5 eq).

In comparison with the limited existing literature for recombinant protein production, the environmental impacts of bench-scale recombinant GFs are substantially larger. When compared to the impacts of simulated commercial-scale ovalbumin production, GF production results in impacts between three to five orders of magnitude larger (Järviö et al., 2021). This may be explained by the differential scales of the production systems (commercial vs bench-scale) as well as the use of variable expression systems (fungus vs bacteria), purification processes (extracellular vs intracellular production), and endpoint applications (egg-white substitute vs biologically active culture media constituent) (Järviö et al., 2021). While reporting lesser impacts, the study on ß-galactosidase provides a better comparison to the present study due to their use of a bacterial host. The GWP and FWE impacts associated with recombinant GF production were six and ten times greater than those associated with ß-galactosidase. The associated impacts on TA and WC were also larger, approximately 4.2 and 1.9 times greater, respectively. SOD was the only impact reduced with IGF-1 production, with a burden 7.6 times smaller than ß-galactosidase (Feijoo et al., 2017).

### 3.2 Global warming potential

The global warming potential (GWP) from the upstream and downstream production of 1 mg of IGF-1 is 0.10 kg CO_2_ eq. As seen in Figure 3, the GWP impacts for the fermentation, cell lysis, and post-elution processes are primarily attributed to electricity usage, whereas in both chromatography processes, GWP impacts are allotted to polypropylene and polyethylene disposable production. Within fermentation, electricity contributes 57% to the process’s GWP, with the second largest contribution attributed to reagent production at 22%. Upon further analysis, tryptone, yeast extract, and sodium chloride within the LB/TB media production sub-process combined contribute to 87% of the fermentation process’s overall reagent impacts. Waste processes are not sizable contributions to fermentation-specific global warming results, contributing to only 1% of the overall process’s impact combined.

**Figure 3.**
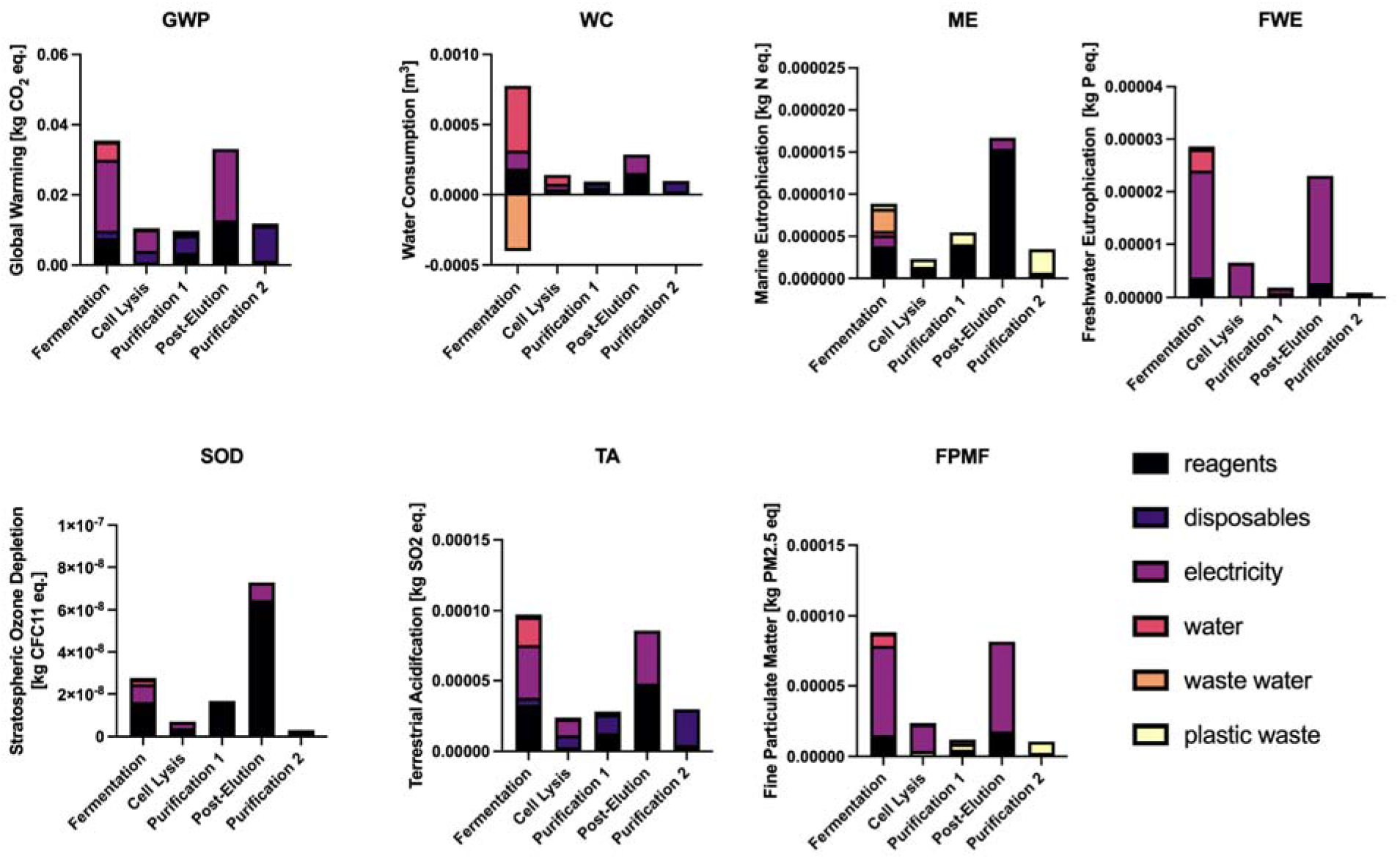
Environmental impact of IGF-1 production per mg. Results and specific contributions by impact category broken down in terms of subsystem 1 (SS1) process (fermentation, cell lysis, first purification, post-elution, or second purification) and input type (reagents, disposables, electricity, water, waste water, or waste plastic).

Further analysis of the sub-processes comprising fermentation reveals that most of the GWP burden associated with electricity usage is ascribed to *E. coli* cultivation, which requires the overnight use of a shaking incubator. As for purification, the post-elution process is the most impactful due to electricity requirements for the overnight mixing and storage of the tagged proteins. Post-elution itself represents 33% of the overall GWP results. As the fusion protein product acquired from the cell lysis process was functionally equivalent to the cleaved, dialyzed protein acquired after post-elution in cell growth studies, elimination of the post-elution step has the potential to lower total GWP to close to half the original GWP results. This scenario is explored further in Section 3.9.

The results of this analysis indicate that the contribution associated with the investigated recombinant GFs to total environmental impacts of CM systems could be significant. While the GF requirements for CM are highly variable, a recent report predicted that the average concentration of IGF-1, FGF, TGF-ß, and PDGF required in future media scenarios could range from 0.0024-0.074 mg/L, with TGF-ß present at the lowest concentration and IGF-1 at the highest (Swartz, 2023). If between 8-42 L of media is needed to produce 1 kg of CM, 0.019-3.11 mg of GF would be necessary (Swartz, 2023). As the GWP of 1 mg GF was estimated to be 0.10-0.18 kg CO_2_ eq at baseline in this study, and the GWP of CM is presently estimated to range from 1.9-26 kg CO_2_ eq (Tuomisto & Teixeira de Mattos, 2011; Tuomisto et al., 2014; Sinke et al., 2023), GFs used in future large-scale production of media could present potentially substantial burdens. GFs are key components in current CM media formulations, with demand estimated to reach 0.6-17.1 tons per year to meet the projected CM demand of 0.4-2.1 million metric tons per year by 2030 (Swartz, 2023). Its impacts indicate the importance of modeling media inputs with concrete processes to accurately estimate the environmental impacts of CM production.

### 3.3 Water consumption

As industrial bioprocesses tend to be water-intensive, water consumption (WC) is a particularly important impact category to consider when assessing the environmental burden of recombinant GF production. The impacts of water usage and reagent production dominated the WC results, accounting for 53% and 42% of the total impacts across all processes, respectively. Within the system, fermentation and post-elution were the most influential on consumptive water usage. For fermentation, the WC impact from reagent, disposable, electricity, and water usage equates to 0.0008 m^3^ per mg of IGF-1 produced. Reagent production and electricity contribute to 25% and 16% of this process’s impacts, respectively, while water has the highest contribution, at 58%. This is due to water usage required to sterilize spent culture medium and to make LB/TB media for fermentation.

The aforementioned results do not include the credit obtained from wastewater treatment. The ReCiPe 2016 Midpoint (H) method attributes negative water consumption values for recycling processes. Spent culture media from cell cultivation as well as wastewater from cleaning processes are assumed to be recycled as average wastewater, providing a credit back to the system that is subtracted from the overall process impacts. In the fermentation process, the treatment of spent media prior to IPTG induction provides a -0.0004 m^3^ credit, decreasing the consumptive water results from this process to 0.0004 m^3^, or 38% of the total system’s WC impacts. Despite the credit warranted from the fermentation process, this process is revealed to still be the primary contributor of the overall consumptive water impacts, with the second purification process closely following at 29% of the total environmental burden. These results can be further mitigated by streamlining the process to only include one purification step.

### 3.4 Marine and Freshwater Eutrophication

The marine eutrophication (ME) burden attributed to IGF-1 was 3.7X10^-5^ kg N eq per 1 mg. The impacts measured for recombinant IGF-1 production can be mostly attributed to the production of the reagents used for the cleavage and dialysis of eluted proteins after purification, resulting in a 45% overall system contribution. Within the cell lysis subunit process as well as the first and second purification subunit processes, waste culminated from plastic disposables accounted for 39% 25%, and 80% of the total marine eutrophication impacts, respectively. As mentioned previously, the removal of the post-elution process enabled by the sufficient functionality of the fusion protein after the first purification process has the potential to decrease ME impacts from both reagent and disposable perspectives.

Conversely, the freshwater eutrophication potential (FWE) attributed to IGF-1 was 6.1X10^-5^ kg P eq per 1 mg. In contrast to marine, freshwater eutrophication impacts are measured through the emission of phosphoric nutrients in aquatic ecosystems. Electricity usage was the most influential input causing the highest impacts across all system processes. This was due primarily to the treatment of spoil from lignite mining within the electricity production process, which caused 63% of the FWE results. As such, it was hypothesized that deviations from baseline electricity usage would appreciably affect impact category outcomes, a hypothesis further explored within the system’s sensitivity analysis (see section 3.7).

### 3.5 Stratospheric ozone depletion and fine particulate matter formation

Per mg IGF-1, the associated stratospheric ozone depletion (SOD) was 1.3X10^-7^ kg CFC11 eq. The dialysis step in the post-elution sub-unit process contributes 57% of the burden as it is a reagent-intensive step that requires 74% of the total glycerol, 73% of the total HEPES, and 40% of the total sodium chloride used in the entire product system.

Reagent use contributes to 82% of the overall burden. As glycerol, the main carbon source, is used in seven of the unit processes within IGF-1 production, it was found to contribute a sizable 83% to reagents’ impact. An average market mix was used for the glycerol input, and thus the impact was found to be largely attributed to the esterification of various oils involved in some of the glycerol production processes included within the mix. However, glycerol is considered a sustainable feedstock option in industry, as it is a renewable, non-edible biomass feedstock that is typically produced as a by-product of biodiesel processes (Dickson et al., 2021). For this reason, future studies may explore the impact minimization associated with the use of a biodiesel by-product as a more accurate representation of the glycerol used in fermentation processes.

The fine particulate matter formation burden (PM) attributed to the production of 1 mg IGF-1 was 2.2X10^-4^ kg PM2.5 eq. Electricity contributed 69% of the PM associated with IGF-1, with energy-intensive processes including fermentation and dialysis being the most impactful. Lignite-, hard coal-, and natural gas-based electricity production carried the heaviest burden within the WECC electricity grid mix used as the baseline for IGF-1 production, which would be expected as combustion-based electricity generation is a known cause of air pollution.

### 3.6 Terrestrial acidification

The TA associated with 1 mg of IGF-1 was 2.6X10^-4^ kg SO_2_ eq. The majority of this burden can be attributed to reagents. Broken down further, 61% of the burden associated with reagents came from glycerol. A substantial portion of the TA burden was associated with electricity and disposables usage as well; 34% and 19% respectively. Production of polyethylene and polypropylene used in plastic tubes and pipettes were the major contributors to disposables impacts, while electricity produced by coal and lignite in the baseline electricity grid mix were the top contributors to the electricity impacts.

### 3.7 Sensitivity analysis of bench-scale IGF-1 results

Glycerol, energy, water, and disposables (polyethylene and polypropylene) were chosen as parameters of interest for sensitivity analysis. These inputs were adjusted to represent a 10% increase and decrease from their baseline values. The model was found to be most sensitive to electricity, with PM and FE being the most extreme. When decreasing electricity usage by 10% from the baseline, PM and FWE impacts decreased 14% and 11%, respectively. Increasing electricity usage led to a 4.5% and 5.7% increase in these impacts. These larger fluctuations associated with changes in electricity use suggested deeper analysis of electricity within the system would be necessary, and thus alternative scenarios were developed and explored in Section 3.8. As the impacts neither increased nor decreased beyond 10% in respect to the baseline for the glycerol, disposables, and water parameters, our results are demonstrated to be robust against 10% fluctuations in this analysis (Figure 4).

**Figure 4.**
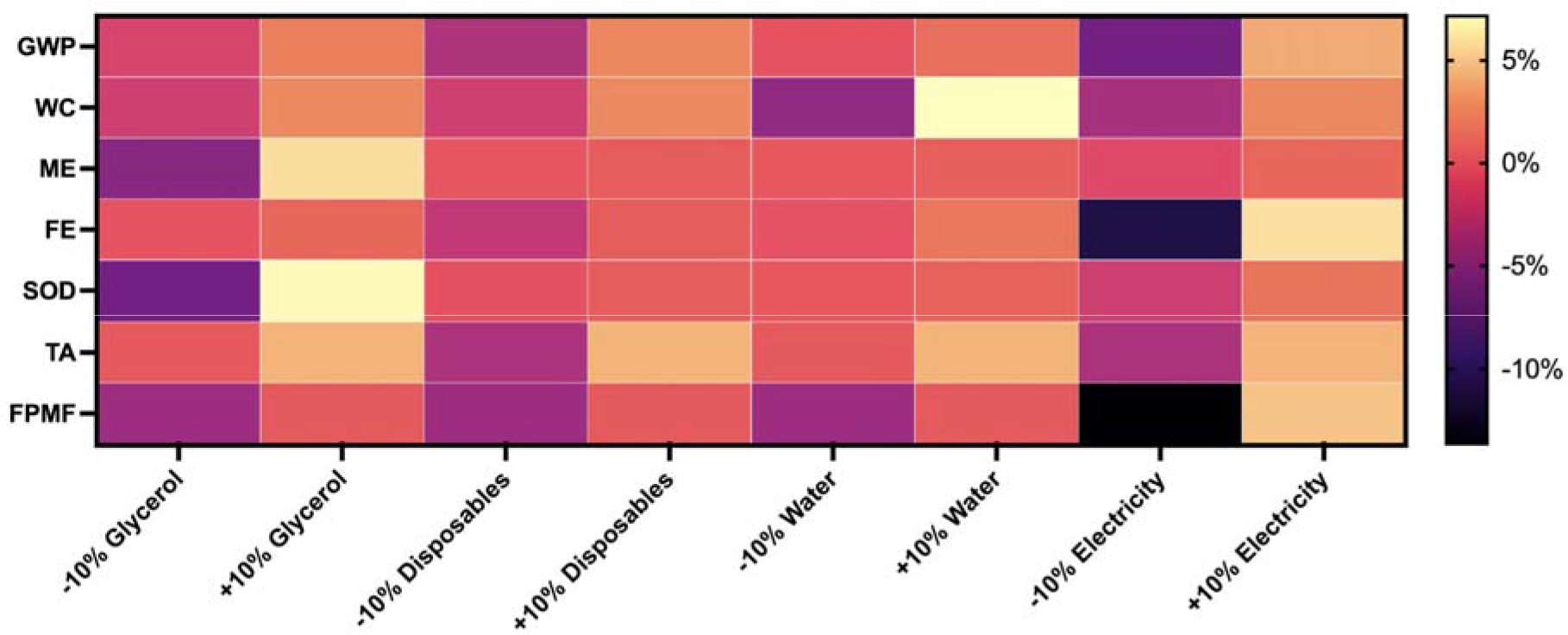
Sensitivity analysis of IGF-1 production. Outcomes for sensitivity analysis represented by heatmap in terms of percent deviation from baseline scenario. Results of analysis based on 10% increase/decrease of inputs within the IGF-1 product system. Inputs include carbon feedstock (glycerol), disposables, water, and electricity use.

### 3.8 Scenario analysis of energy use

Due to findings from the sensitivity analysis, scenarios were developed to model alternative electrical transmission grids to understand the effects of regional specific power sources on the results. The alternative US scenarios chosen were the MRO and NPCC electricity grid mixes, which include varying proportions of renewable and nuclear energy sources used to produce electricity (supplementary material). Comparisons between both major electricity grids, the Eastern Interconnection and the Western Interconnection (WECC), the latter which serves as the model’s baseline, allows for a holistic comparison of electricity scenarios across the entire US.

Analysis of the IGF-1 production system using the MRO electricity grid mix resulted in similar or increased environmental burdens across all categories relative to baseline (Figure 5). As such, the MRO regional electricity scenario represents the worst performance for electricity impacts of the scenarios explored, as it relies on the lowest amounts of renewable and nuclear sources (supplementary material). The NPCC grid mix resulted in appreciable decreases from the baseline WECC mix in terms of PM at -50% and GWP at -18%. FE was the most heavily impacted by changes in electricity grids. The Midwest (MRO) scenario resulted in an 82% increase in this impact category, while the Northeast scenario (NPCC) resulted in a 67% decrease. FE linked with electricity use was driven by use of lignite coal for electricity generation. Similarly impacted was PM, TA, and GWP of the product system. Use of electricity from the MRO grid resulted in a 62% increase in kg PM2.5 eq, a 34% increase in kg SO_2_ eq, and a 20% increase in CO_2_ eq due to the relatively large portion of electricity sourced from lignite and coal. The NPCC grid resulted in a 50% decrease in PM and an 18% decrease in GWP.

**Figure 5.**
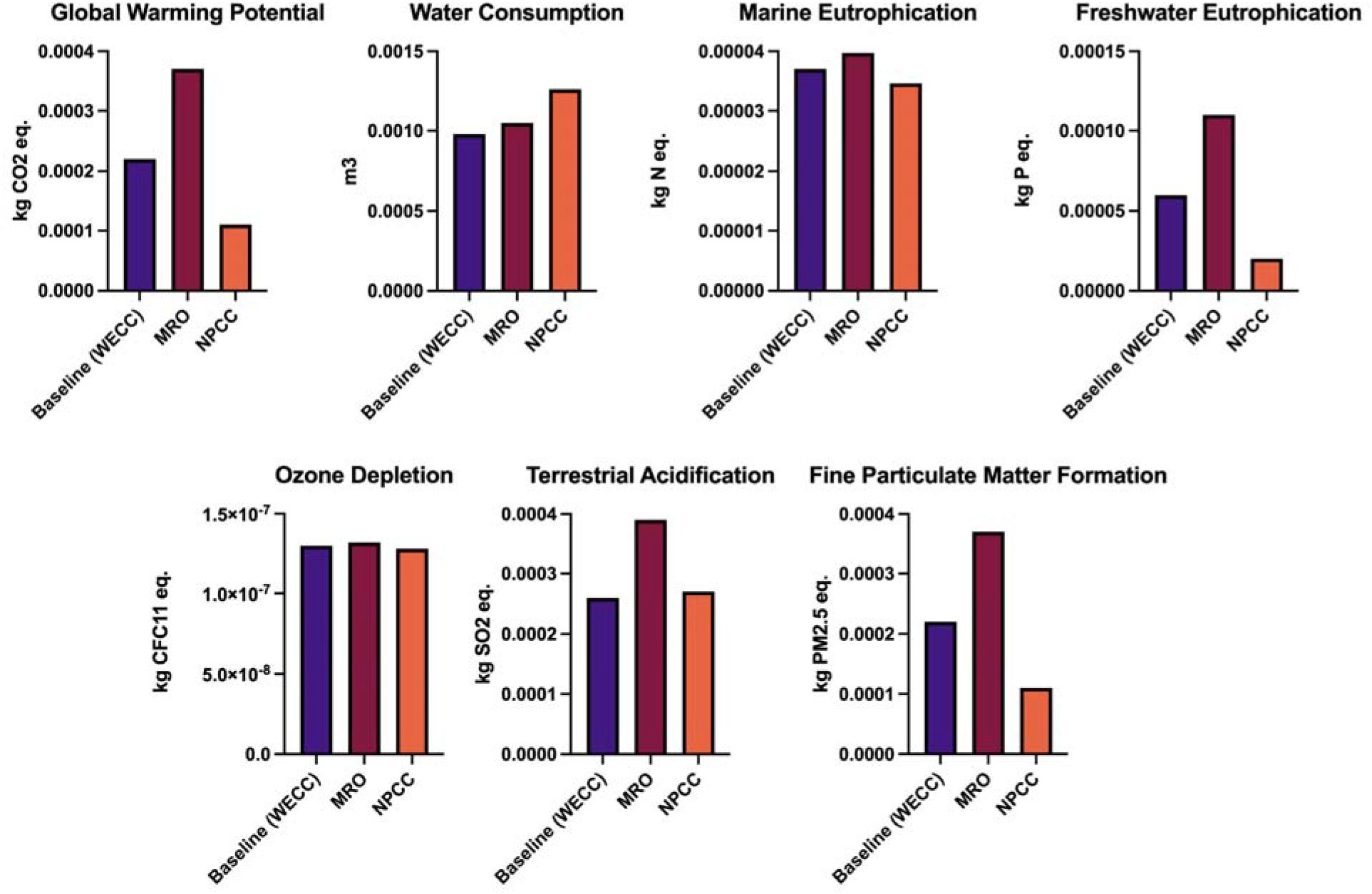
Comparison of environmental impacts of IGF-1 production scenarios using different electricity grid scenarios. Results for IGF-1 production by impact category across baseline (WECC) MRO and NPCC electricity grids.

### 3.9 Scenario analysis for process optimization

The functionality of the fusion protein following the first IMAC purification process was confirmed through additional cell growth studies performed by the collaborating laboratory (Venkatesan et al., 2022). Therefore, following elution, the protein sample may be directly dialyzed in a storage buffer, concentrated, and flash frozen. With the elimination of the overnight steps, the impacts across all impact assessment categories are seen to decrease by an average of 46%, with the largest decrease apparent for FE at 64%. Similarly large decreases in SOD and ME were observed at 61% and 60% respectively (Figure 6). These results further reveal the importance of product-specific recombinant protein production LCA for accurate CM environmental assessments. Whereas recombinant proteins necessary for pharmaceutical purposes require purity at the highest percentage, recombinant proteins utilized for food applications not necessarily present in the final food product may not require high purity for functionality (Waschulin & Specht, 2018). The importance of optimization of such processes to lower environmental impacts is justified.

**Figure 6.**
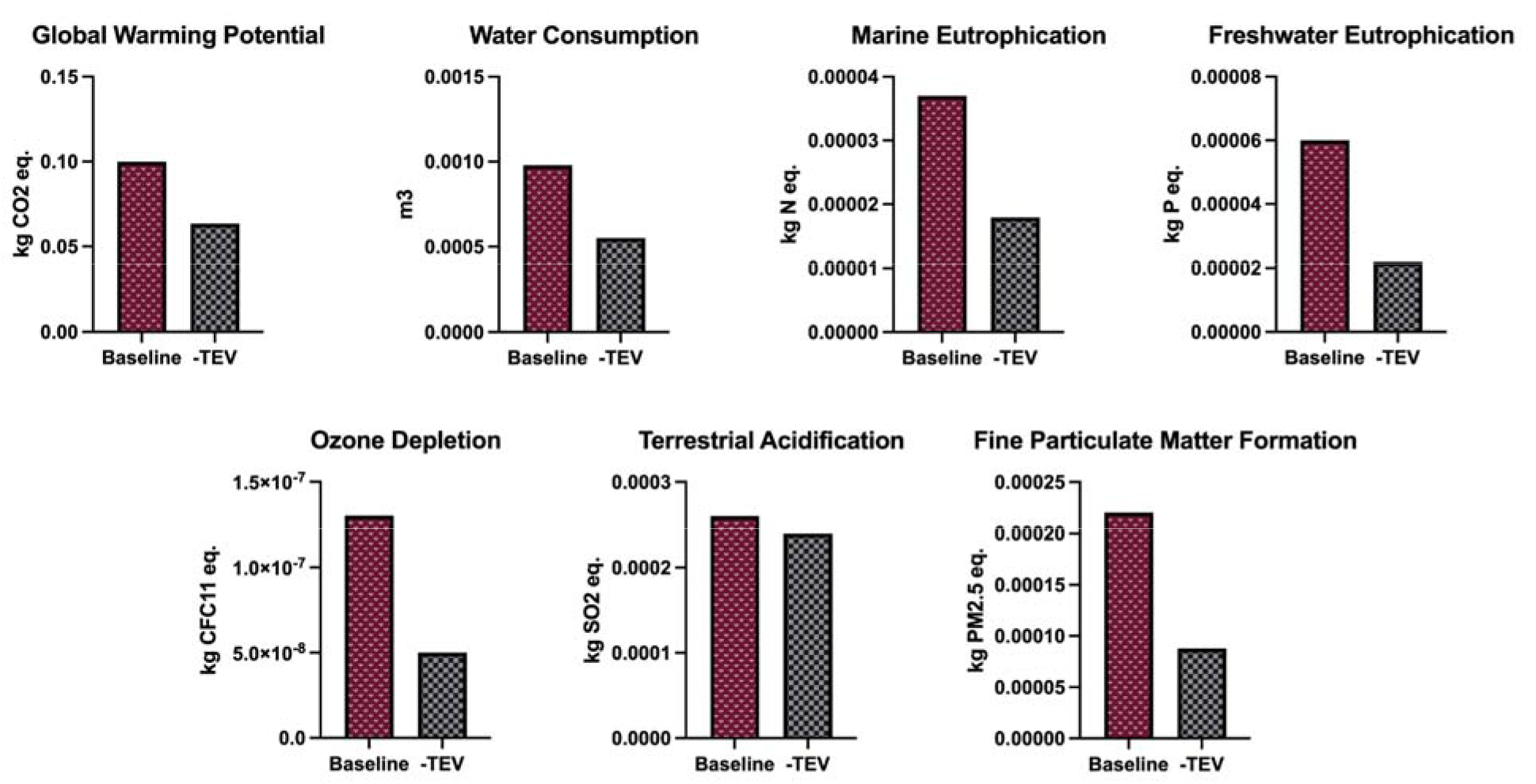
Comparison of environmental impacts of IGF-1 production scenarios considering a streamlined process. Results for IGF-1 production by impact category for baseline (+TEV) and optimized process (–TEV) scenarios.

### 3.10 Limitations of the study

This LCA study quantified the life cycle environmental impacts of four GFs developed by a procedure suggested by Venkatesan et al. (2022). However, the environmental impacts of GF production might be varied if other procedures are used for development. Further research should investigate the environmental impacts of plant-derived GFs, as 79% of suppliers and 67% of manufacturers in the industry are seeking ways to produce growth factors from non-mammalian sources (Swartz, 2021). Another limitation of this work is the scale of the analysis. It is suggested that only future bench scale LCA studies apply the inventories developed in this study as the environmental impacts of industrial scale GFs may be lower, assuming ecological efficiencies are achieved at larger scale. Since the CM industry is in its earliest stages of development, we expect bench-scale environmental analyses of CM-based products to proliferate in the near future, particularly those focused on benchmarking and process optimization. Given the potential influence of GFs on environmental performance of CM production systems, these studies could benefit from integrating and adapting the inventory data from this study.

Uncertainty of results is a challenging issue in every LCA study due to uncertainties from, among others, the accuracy and scope of the collected data, model assumptions, functional unit, and impact assessment selection (Nikkhah et al., 2021). Likewise, in this study, several assumptions had to be made to estimate the necessary data for the LCA. A comprehensive list of assumptions and the processes and proxies used to develop the LCI are provided in Supplementary Material 2 and 3.

## 5. Conclusions

The development of a publicly-available GF LCI and LCA has critical applications for future studies aimed to quantify the environmental impacts of CM production. The present study collected and assessed the inputs and outputs of a novel bench-scale recombinant GF system to develop LCIs for four recombinant GFs (IGF-1, FGF, TGF-ß, and PDGF). Analysis of the system using ReCiPe 2016 Midpoint (H) LCIA methods for GWP, WC, FWE, ME, SOD, TA, and PM impact categories found high burdens that were driven largely by electricity and substrate consumption. Uncertainty was accounted for through sensitivity analysis of high impact inputs, specifically glycerol, electricity, water, and disposable usage, as well as scenario analysis of alternate electricity grids and production process optimization. The results of the sensitivity and scenario analyses demonstrated the robustness of the results to input fluctuations, with the exception of electricity. Sensitivity in electricity burdens were found to be impact category-dependent, with the sources of electricity generation causing most of the variation in outcomes. Overall, when comparing the results of this analysis against published CM LCAs, the impacts of GF production could be substantial, falling between 0.38% - 9.5% of total GWP impacts.

Bench-scale processes generally yield low products relative to their necessary inputs, and thus increasing efficiency through scale is critical in biotechnological processes. For use in further CM LCA studies, it is important to not only model the recombinant production of GFs, but to also model this system at the correct scale and degree of purity. Industrially, these proteins are produced at the scale and grade of pharmaceuticals and are therefore identified as a hotspot in both economic and environmental analyses. It is hypothesized that if GFs were produced for the purpose of food production, which entails a larger scale and lower standards of purity, their impacts may prove to be meaningfully lower. As such, ex-ante (i.e., forecast the impact of a product before its industrial production) LCA studies should be conducted at the appropriate scale to capture economies of scale for GF production by performing an analysis using integrated methods to simulate scale-up of the process.

## Author Contributions

Conceptualization, K.T., R.A., and N.T.B.; methodology, K.T., R.A., C.C., N.T.B.; software, R.A., K.T., A.N., and N.T.B.; validation, K.T., R.A., A.N., C.C., N.T.B.; formal analysis, K.T., R.A., A.N., N.T.B.; resources, R.A., K.T., and C.S.; data curation, C.S., R.A., and K.T.; writing—original draft preparation, K.T., R.A.; writing—review and editing, K.T., R.A., A.N., C.C., C.S., N.T.B., and D.K.; visualization, K.T., R.A.; supervision, N.T.B.; funding acquisition, K.T., R.A., N.T.B., and D.K. All authors have read and agreed to the published version of the manuscript.

## Data Availability Statement

The raw data inventories, the processes and assumptions used for the LCA, and the life cycle inventories of emissions are provided in the Supplementary Material.

## Acknowledgments

This work was supported by the Agriculture and Food Research Initiative (AFRI) Sustainable Agricultural Systems program, grant no. 2021-699012-35978 from the USDA National Institute of Food and Agriculture. K.T. and R.A. were also supported by the Tufts Institute for the Environment Fellowship. C.S. was supported by the New Harvest Doctoral Research Fellowship.

## Conflicts of Interest

The authors declare no conflicts of interest.

**Appendix 1.**
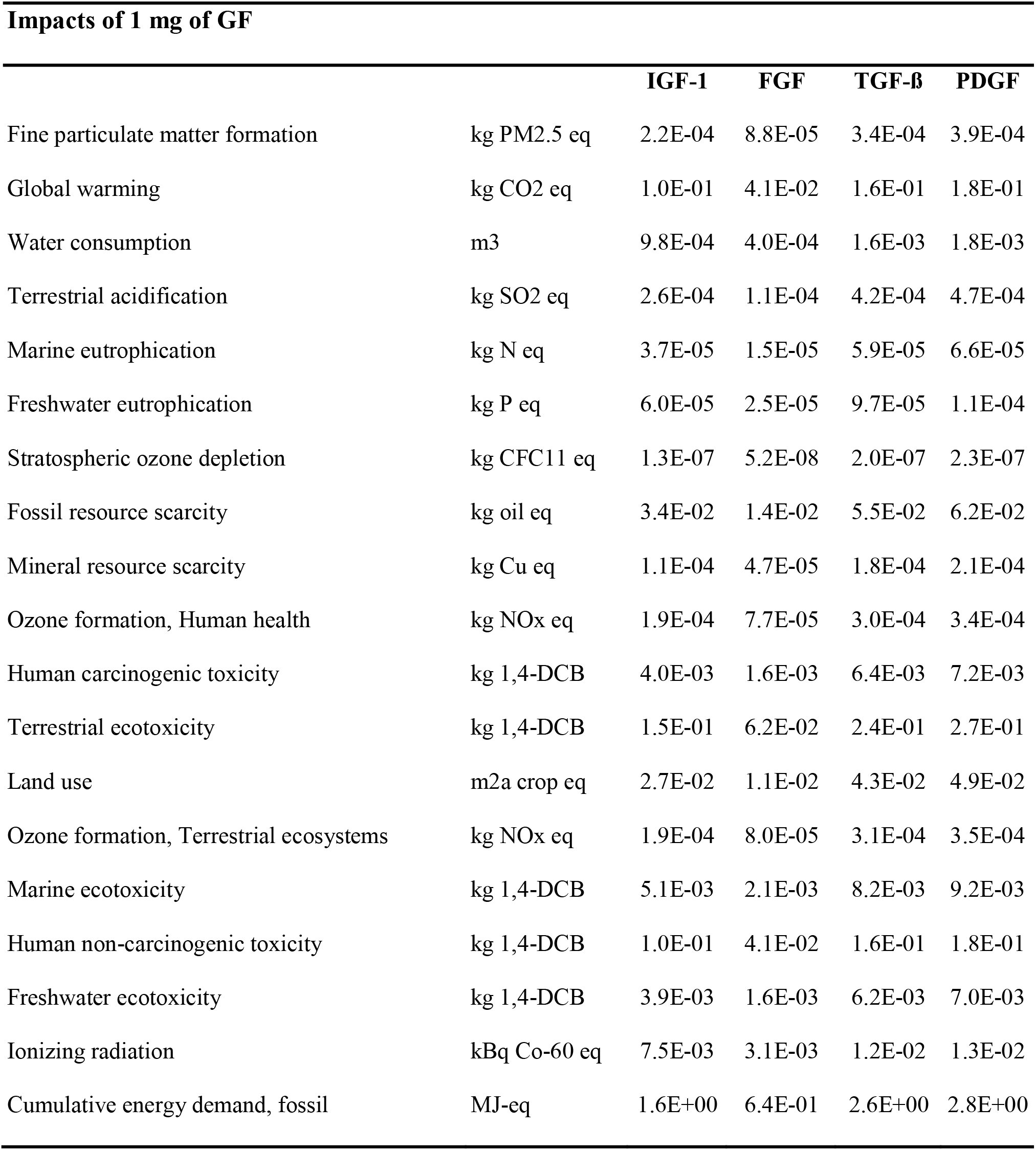
Complete ReCiPe 2016 Midpoint (H) results for IGF-1, FGF, TGF-ß, and PDGF product systems.

---

**Supplementary Table 1.1.**
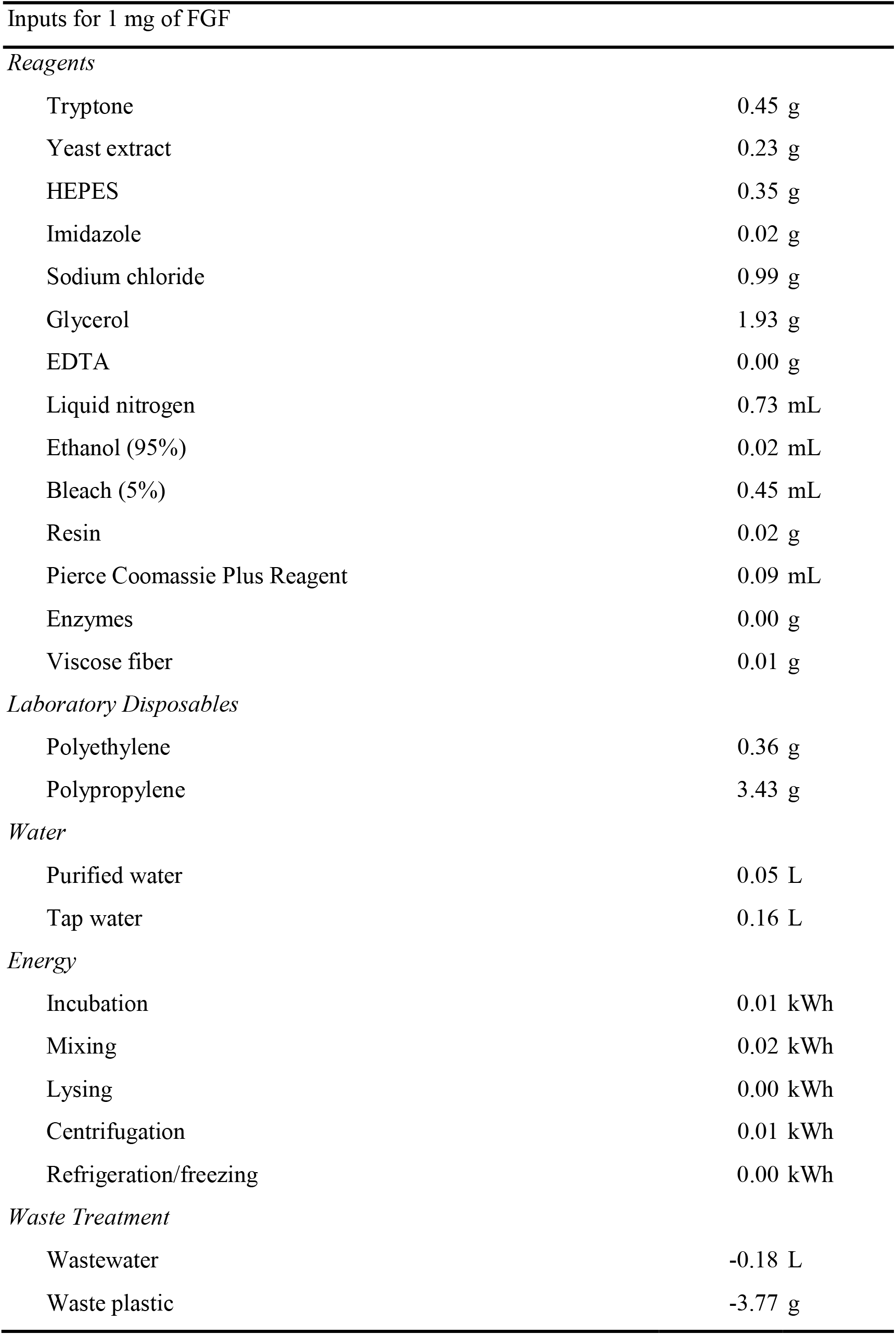
Inventory summary of bench-scale FGF (FU=1 mg FGF)

|  |  |
| --- | --- |
| Tryptone | 0.45 g |
| Yeast extract | 0.23 g |
| HEPES | 0.35 g |
| Imidazole | 0.02 g |
| Sodium chloride | 0.99 g |
| Glycerol | 1.93 g |
| EDTA | 0.00 g |
| Liquid nitrogen | 0.73 mL |
| Ethanol (95%) | 0.02 mL |
| Bleach (5%) | 0.45 mL |
| Resin | 0.02 g |
| Pierce Coomassie Plus Reagent | 0.09 mL |
| Enzymes | 0.00 g |
| Viscose fiber | 0.01 g |
| *Laboratory Disposables* |  |
| Polyethylene | 0.36 g |
| Polypropylene | 3.43 g |
| *Water* |  |
| Purified water | 0.05 L |
| Tap water | 0.16 L |
| *Energy* |  |
| Incubation | 0.01 kWh |
| Mixing | 0.02 kWh |
| Lysing | 0.00 kWh |
| Centrifugation | 0.01 kWh |
| Refrigeration/freezing | 0.00 kWh |
| Wastewater | -0.18 L |
| Waste plastic | -3.77 g |

**Supplementary Table 1.2.**
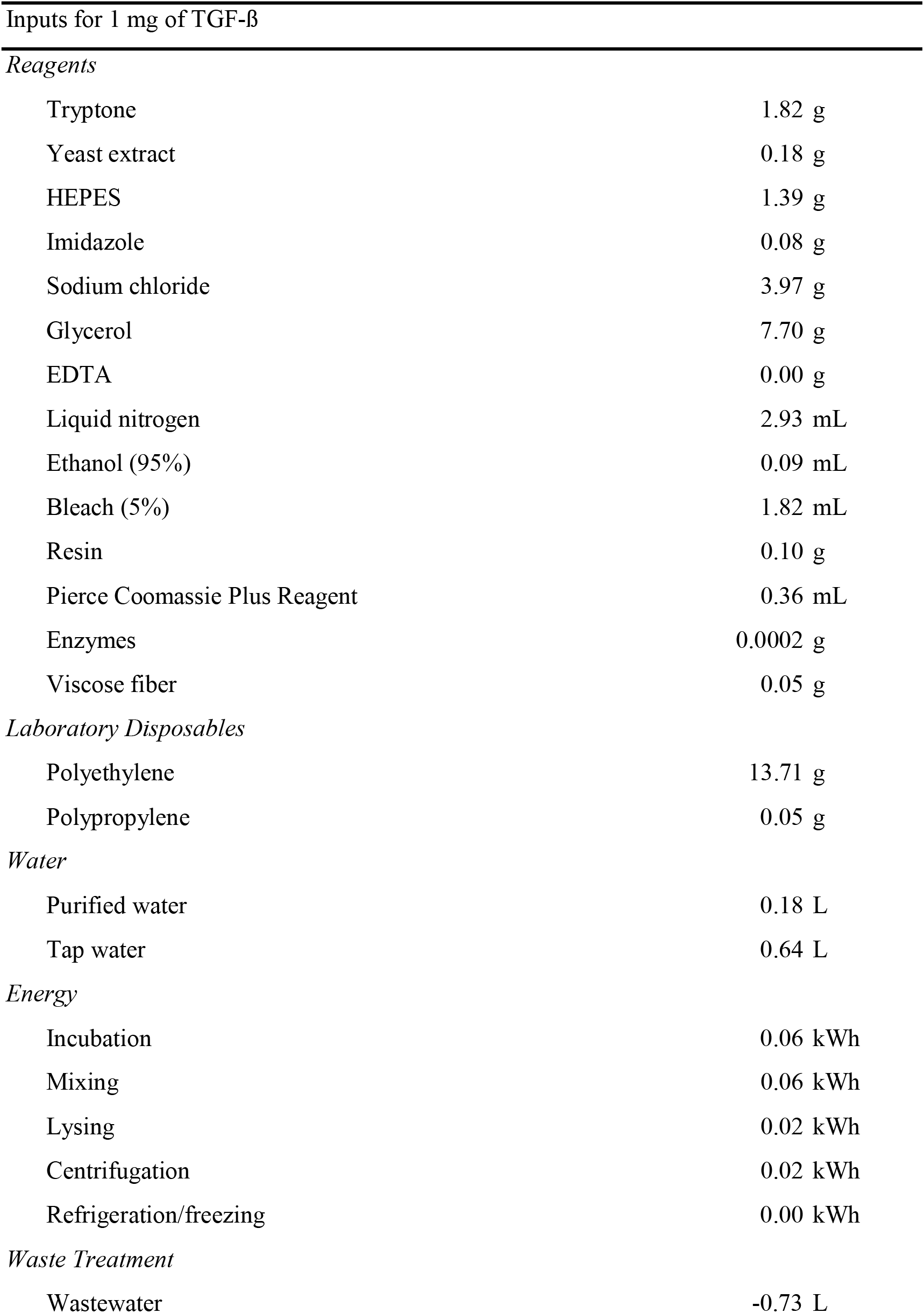
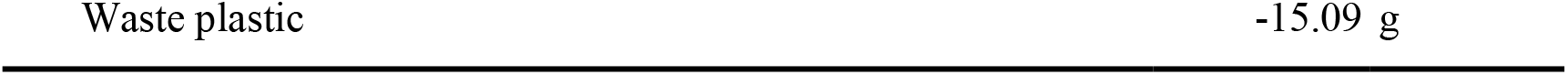
Inventory summary of bench-scale TGF-ß (FU=1 mg TGF-ß)

**Supplementary Table 1.3.**
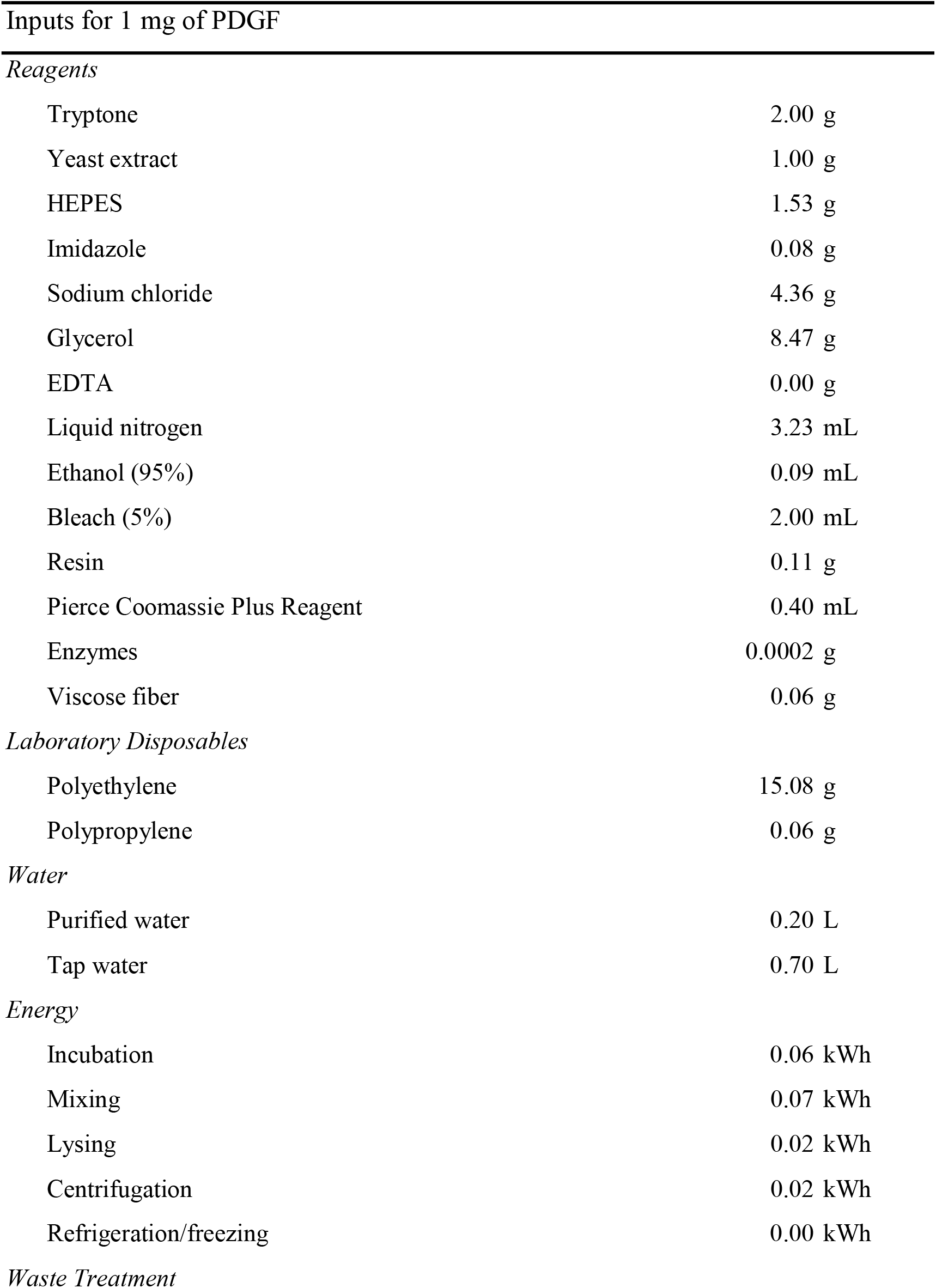
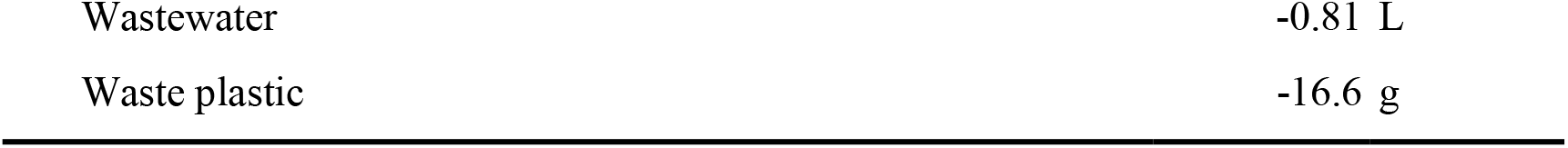
Inventory summary of bench-scale PDGF (FU=1 mg PDGF) Inputs for 1 mg of PDGF

|  |  |
| --- | --- |
| Tryptone | 2.00 g |
| Yeast extract | 1.00 g |
| HEPES | 1.53 g |
| Imidazole | 0.08 g |
| Sodium chloride | 4.36 g |
| Glycerol | 8.47 g |
| EDTA | 0.00 g |
| Liquid nitrogen | 3.23 mL |
| Ethanol (95%) | 0.09 mL |
| Bleach (5%) | 2.00 mL |
| Resin | 0.11 g |
| Pierce Coomassie Plus Reagent | 0.40 mL |
| Enzymes | 0.0002 g |
| Viscose fiber | 0.06 g |
| *Laboratory Disposables* |  |
| Polyethylene | 15.08 g |
| Polypropylene | 0.06 g |
| *Water* |  |
| Purified water | 0.20 L |
| Tap water | 0.70 L |
| *Energy* |  |
| Incubation | 0.06 kWh |
| Mixing | 0.07 kWh |
| Lysing | 0.02 kWh |
| Centrifugation | 0.02 kWh |
| Refrigeration/freezing | 0.00 kWh |
| *Waste Treatment* |  |
| Wastewater | -0.81 L |
| Waste plastic | -16.6 g |

**Supplementary Material 2.**
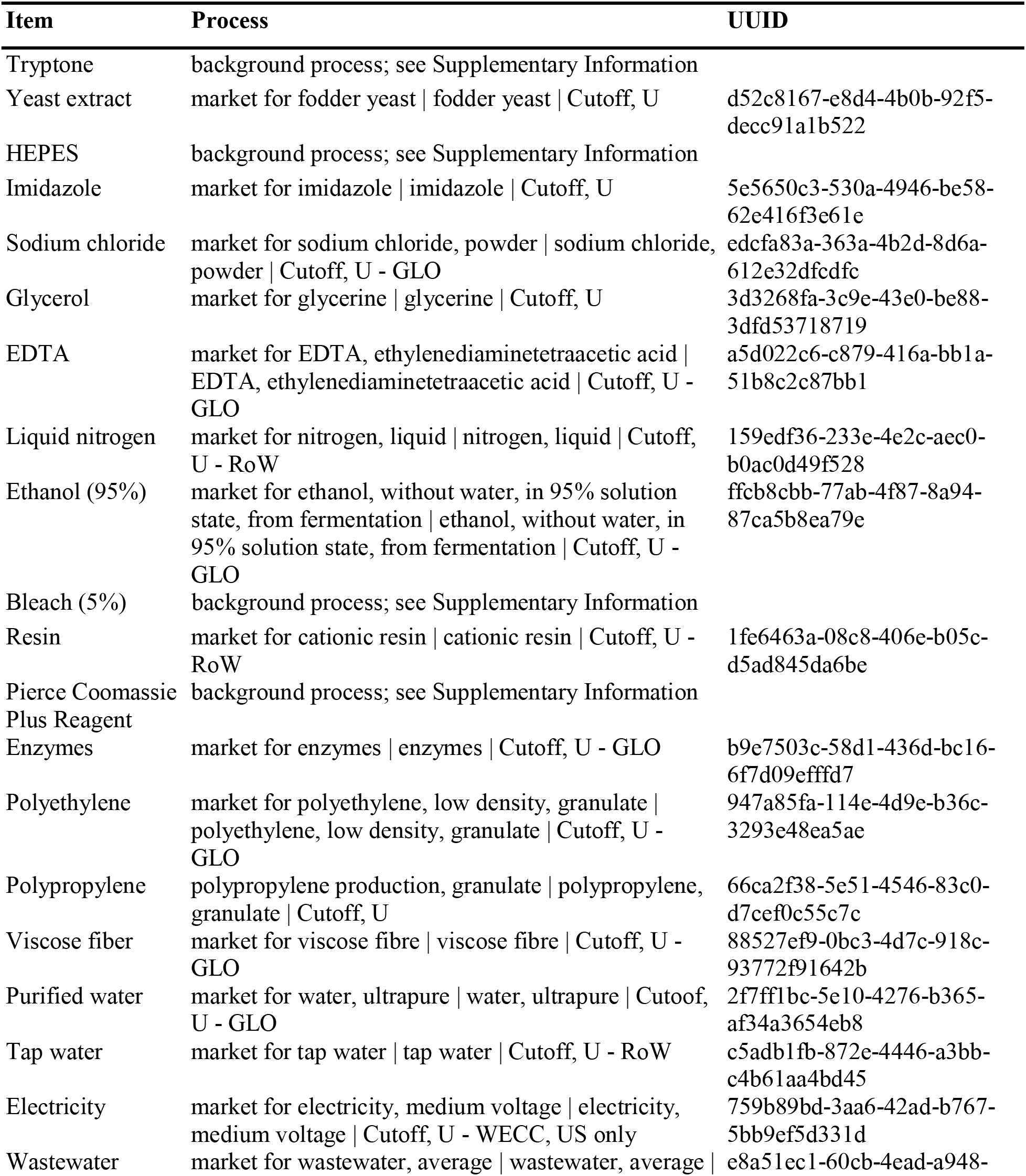
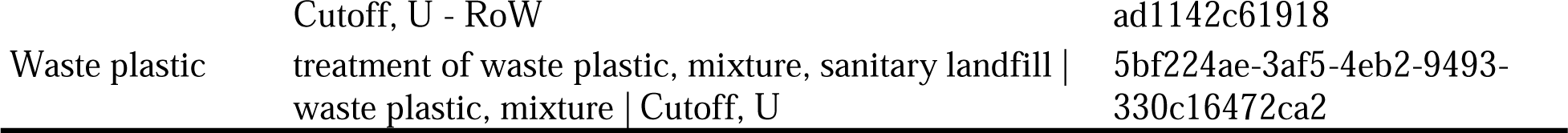
Processes used for LCA inventory development for growth factor production

**Supplementary Material 3.**
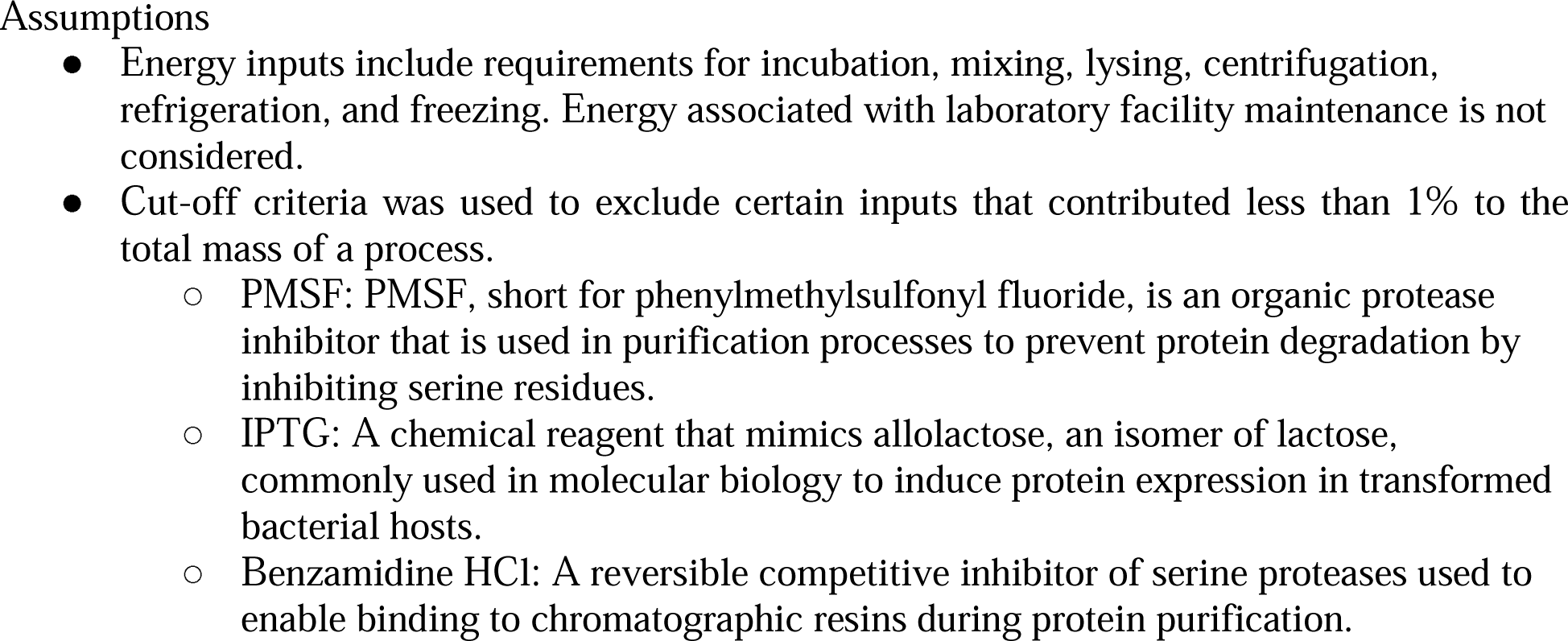
Assumptions made for data collection and the LCA analysis

**Supplementary Table 3.1.**
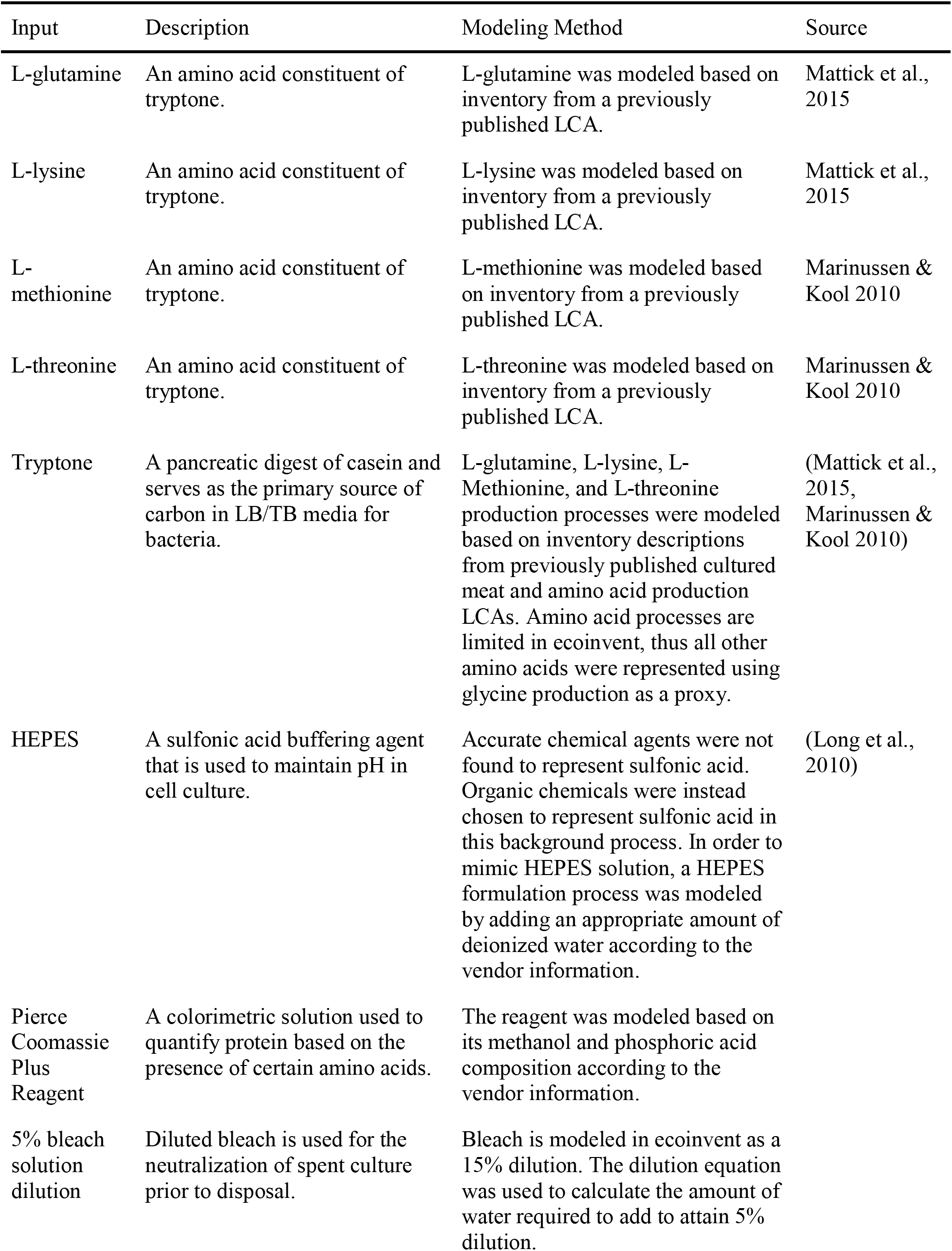
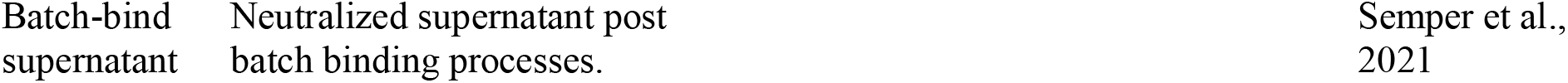
Background processes modeled in ecoinvent.

**Supplementary Table 3.2.** Proxies lacking an exact or near match in ecoinvent and excluded

| Input | Description | Proxy/Cutoff | Rationale |
| --- | --- | --- | --- |
| Yeast Extract | Used in cell culture as a media additive to provide cells with necessary nutrients such as amino acids, peptides, vitamins, and carbohydrates. | Fodder yeast | Fodder yeast flow was used in multiple processes for the production of protein and energy feeds. |
| HisPur™ Ni-NTA Resin | Nickel-IMAC resin used for the purification of His-tagged fusion proteins composed of nickel-charged nitrilotriacetic acid (NTA) on microscopic 6% cross linked agarose beads. | Cationic resin | Due to the microscopic nature of the agarose beads, nitrilotriacetic acid was chosen as a proxy to represent the resin. |
| TEV Protease | Enzyme from tobacco etch virus (TEV) used for the cleavage of fusion proteins and removal of recombinant protein purification tags | Enzymes | Generic enzymes were assumed to be accurate for this input. |
| SnakeSkin Dialysis Tubing | Disposable input used in dialysis composed of regenerated cellulose material. | Viscose fibre | Viscose fibre was chosen as it is a type of regenerated cellulose material. |
| PMSF | An organic protease inhibitor that is used in purification processes to prevent protein degradation by inhibiting serine residues. | Cutoff | <1% mass contribution to the overall |
| IPTG |  | Cutoff |  |
| Benzamidine HCl |  | Cutoff |  |

